# DeepSea: An efficient deep learning model for single-cell segmentation and tracking of time-lapse microscopy images

**DOI:** 10.1101/2021.03.10.434806

**Authors:** Abolfazl Zargari, Gerrald A. Lodewijk, Najmeh Mashhadi, Nathan Cook, Celine W. Neudorf, Kimiasadat Araghbidikashani, Stefany Rubio, Eva Hrabeta-Robinson, Angela N. Brooks, Lindsay Hinck, S. Ali Shariati

## Abstract

Dynamics and non-genetic heterogeneity are two fundamental characteristics of basic processes of life such as cell division or differentiation. Time-lapse microscopy is the only method that can directly capture the dynamics and heterogeneity of fundamental cellular processes at the singlecell level with high temporal resolution. Successful application of single-cell time-lapse microscopy requires automated segmentation and tracking of hundreds of individual cells over several time points. Recently, deep learning models have ushered in a new era in the quantitative analysis of microscopy images. However, integrated segmentation and tracking of single cells remain challenges for the analysis of time-lapse microscopy images. This work presents a versatile and trainable deep-learning software, termed DeepSea, that allows for both segmentation and tracking of single cells in sequences of phase-contrast live microscopy images. Our segmentation model can easily be trained to segment phase-contrast images of different cell types with higher precision than existing models. Our tracking model allows for quantification of dynamics of several cell biological features of individual cells, such as cell division cycle, mitosis, cell morphology, and cell size, with high precision using phase-contrast images. We showcase the application of DeepSea by analyzing cell size regulation in embryonic stem cells. Our findings show that embryonic stem cells exhibit cell size control in the G1 phase of the cell cycle despite their unusual fast division cycle. Our training dataset, user-friendly software, and code are available here https://deepseas.org.

## Introduction

Cells are frequently adapting their behavior in response to environmental cues to make important fate decisions, such as whether to divide or not. In addition, individual cells within a clonal population and under identical conditions display heterogeneity in response to environmental cues [1]. In recent years, it has become clear that single-cell level analysis over time is essential for revealing the dynamics and heterogeneity of individual cells [2, 3].

Single-cell quantitative live microscopy can directly capture both dynamics and heterogeneity of cellular decisions by continuous long-term measurements of cellular features [4, 5]. Widely available microscopy techniques such as label-free phase-contrast live microscopy allow for monitoring the dynamics of morphological features such as the size and shape of the cells [6]. The key to the successful application of single-cell live microscopy is the scalable and automated analysis of a large dataset of images. Typical live-cell imaging of biological features of cells is a multi-day experiment that produces several gigabytes of images collected from thousands of cells [7]. A major challenge for quantitative analysis of these images is the difficulty of accurately defining the borders of a cell, segmentation, and tracking them over time. Low signal-to-noise ratio, existing non-cell small particles in the background, the close proximity of cells, and unpredictable movements are among the challenges for software-based automated image analysis of live single-cell microscopy data. In addition, cells are non-rigid bodies, and thus, tracking them is more challenging because they can change their shapes with time. Most critically, they divide into two new daughter cells during mitosis, which is unique and not comparable with other phenomena we encounter in conventional object tracking applications. Solving single-cell microscopy challenges requires integrating different disciplines, such as cell biology, image processing, and machine learning.

In recent years, deep learning (DL) has outperformed conventional rule-based image processing techniques in tasks such as object segmentation and object tracking [8–10]. Traditional image segmentation approaches often require experiment-specific parameter tuning, while DL schemes are adaptive and trainable. More recently, DL-based image processing methods have attracted attention among cell biologists and microscopists, for example, to: localize single molecules in super-resolution microscopy [11], enhance the resolution of fluorescence microscopy images [12], develop an automated neurite segmentation system using a large 3D anisotropic electron microscopy image dataset [13], design a model to restore a wide range of fluorescence microscopy data [14], and train a fast model that refocuses 2D fluorescence images onto 3D surfaces within the sample [15]. In particular, DL-based segmentation methods have greatly facilitated the task of cell body segmentation in microscopy images [16–19]. However, the successful application of DL-based models for time-lapse microscopy depends on the integration of segmentation and tracking in one platform to automate the analysis of a large sequence of images.

Here, we developed a versatile and trainable deep learning model for cell body segmentation and cell tracking in time-lapse phase-contrast microscopy images of mammalian cells at the singlecell level. Using this model, we developed a user-friendly software, termed DeepSea, for automated and quantitative analysis of phase-contrast microscopy images. We showed that DeepSea is able to capture dynamics and heterogeneity of cellular features such as cell cycle division and cell size in different cell types. Our analysis of cell size distribution in mouse embryonic stem cells revealed that despite their short G1 phase of the cell cycle, embryonic stem cells exhibit cell size control in the G1 phase of the cell cycle.

## Results

### Training a cell segmentation and tracking model

An overview of our approach is illustrated in Figure 1. We generated and collected a dataset of image sequences of three cell types, including 1) mouse embryonic stem cells (1174 images), 2) bronchial epithelial cells (2010 images), and 3) mouse C2C12 muscle progenitor cells (502 images). Individual cells were manually annotated and labeled to generate a training dataset, including pairs of original cell images and corresponding cell ground-truth mask images, using our in-house Matlab-based cell segmentation software (available here https://deepseas.org/software/). Next, we used the annotated dataset of cell images to train a supervised DL-based segmentation model called DeepSea to detect and segment the cell bodies (Figure 1A,B). To be able to visualize the dynamics of cellular behavior, we added cell tracking capability to our DeepSea model (Figure 1C). We trained a DL tracking model involving two separate input paths fed by the segmented cell images of times t-τ and t. With this model, we could monitor multiple cellular phenotypes and several cell division cycles across the microscopy image sequences and generate lineage tree structures of cells. To make our model widely accessible, we developed a DL-based software with a graphical user interface that allows researchers with no background in machine learning to automate the measurement of cellular features of live microscopy data (Figure 1D). We added manual editing options to DeepSea to allow users to correct our model outputs when needed. In addition, our software allows researchers to train a new model with a new cell type annotated dataset. We provide step-by-step instruction on how to use our software and train a model with a new dataset. Our software, instructions, and cell images dataset are publicly available on https://deepseas.org/.

**Figure 1:**
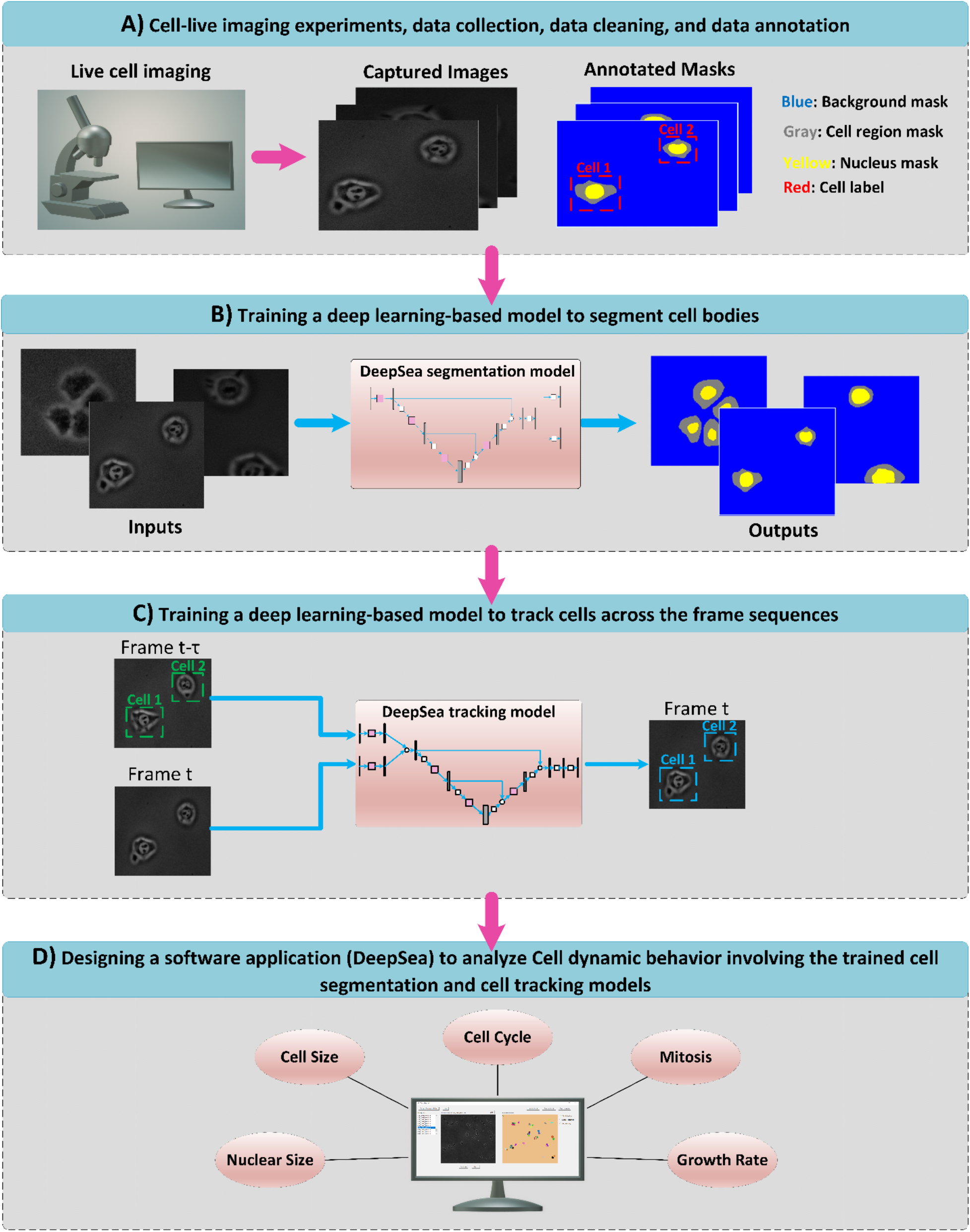
An overview of our approach to collect microscopy data (A), design deep learning models for segmentation (B) and tracking (C) and development of a user-friendly software(D) to analyze cell biological features in live microscopy data.

### DeepSea performance evaluation

After successful training of the DeepSea model, we evaluated our model’s segmentation and tracking performance using a set of test images. The trained segmentation model fits the exact boundary of the target cells and labels their pixels with different colors, helping to determine each cell’s shape and area within the input microscopy image (Supplementary Figure 1). To evaluate the performance of our segmentation model, we matched the model’s predictions to true manually segmented cell bodies at different thresholds of the standard intersection over union metric (IoU) on the test images. Next, we used the standard average precision metric which is commonly used in pixel-wise segmentation and object detection tasks, to compare DeepSea with two recently developed segmentation models [16, 17]. DeepSea is able to outperform existing state-of-the-art models such as CellPose [16] and StarDist [17] in terms of latency and mean average precision (mAP) when trained on the same training sets and tested on the same test sets at all IoU thresholds (Figure 2A,B). Notably, we observed similar prediction accuracy between images with a higher density of cells with touching edges (hard cases) and images with a lower density of cells (easy) with an overall higher precision compared to the CellPose model (Figures 2C,D). In addition, we demonstrated the generalizability of the DeepSea model performance with different cell type test images (Figure 2E). Three examples of the DeepSea and CellPose segmentation model’s output are compared in Figure 3. We also showed that DeepSea is accurate with high-density cell cultures (Supplementary Figure 2). Together, these results indicate that DeepSea’s segmentation model works robustly across different densities of cells and different cell types with high prediction accuracy.

**Figure 2:**
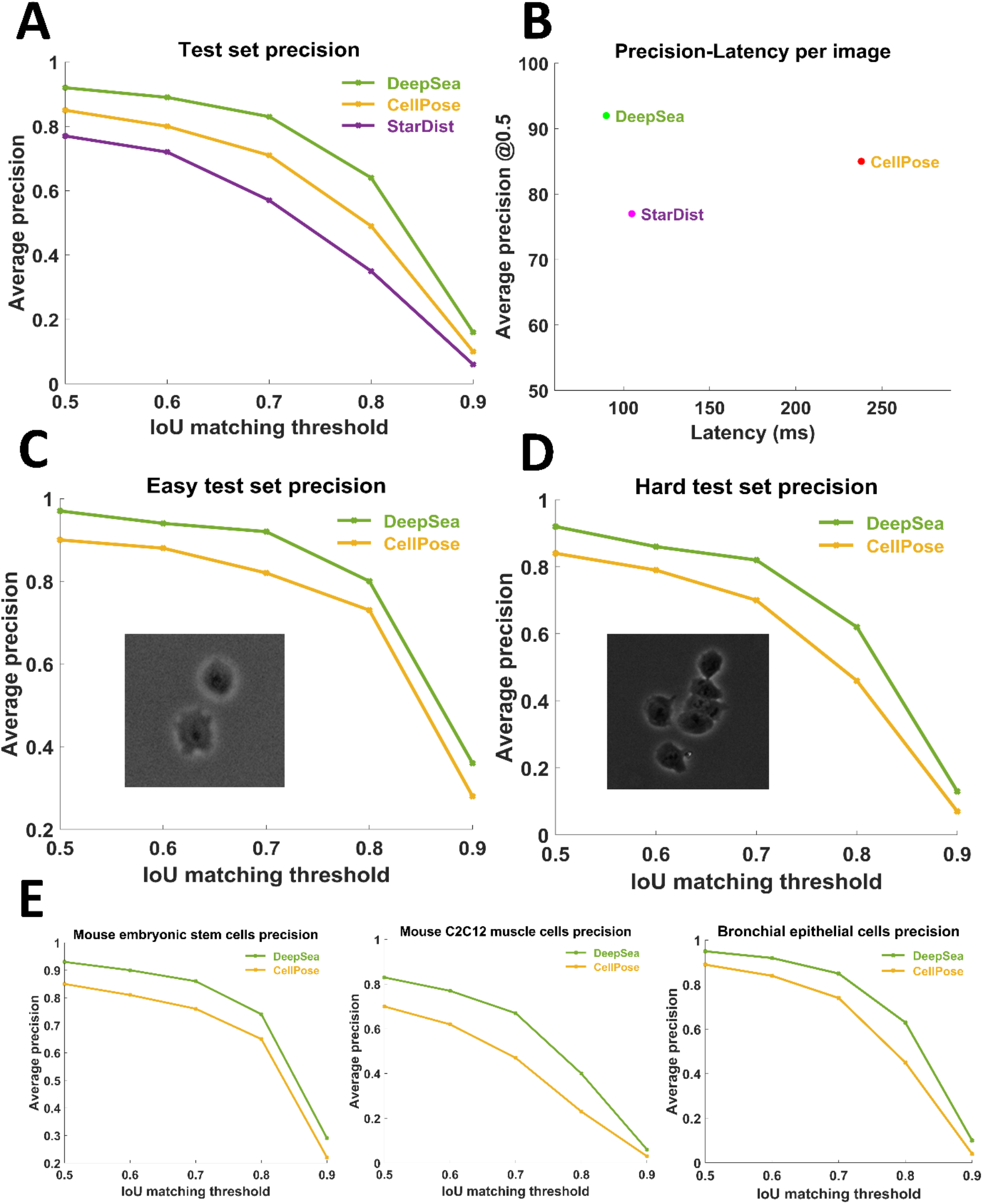
Segmentation model evaluation on the test set images. **A**) Comparing the performance of DeepSea, Cell Pose, and StarDist using the standard average precision at different IoU matching thresholds. **B**) Measuring models’ latency (per image) to compare the DeepSea efficiency with the other models. **C-D**) Comparing models’ performance in segmenting easy (sparse cell density) and hard test cases (high cell density). **E**) Comparing models’ performance in segmenting different cell types.

**Figure 3).**
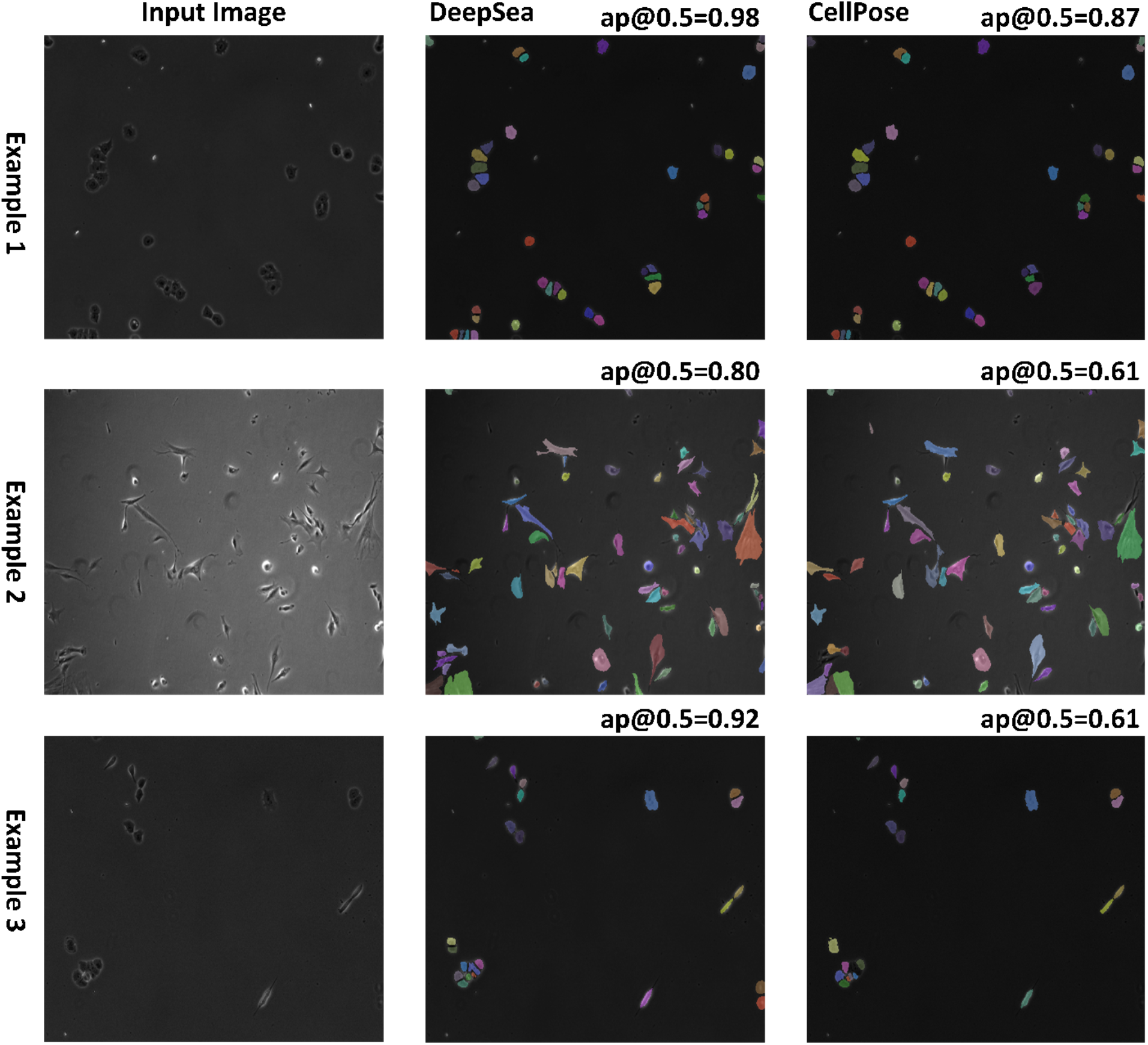
Three examples of DeepSea output (middle column) compared with the CellPose (left column) for different cell types are shown. DeepSea has higher average precision when compared to CellPose (ap).

DeepSea tracking model inputs the target cell image at the previous time point and the cell image at the current time point to generate a binary cell body mask of the target cell at the current time point (Supplementary Figures 3-4). For the tracking model, we evaluated the model’s performance on the test set by measuring the average precision of single-cell tracking from one frame to the next frame as well as mitosis detections. We matched the target cell body masks obtained from the tracking model at time t to the true target cell bodies (at time t) at different matching thresholds of IoU. While our model achieved 99% precision (@0.5 IoU threshold) for tracking single cells, the precision of our model for mitosis detection was around 89% (@0.5 IoU threshold) (Figure 4A,B). Mitosis detection was particularly challenging for stem cells (Supplementary Figure 5). There is likely a direct relationship between the mitosis detection results and the frequency of imaging intervals (aka sampling rate) as the bronchial epithelial cell images were sampled every 5 minutes, while most stem cell samples’ sampling rate is 30 min. More frequent imaging of microscopy frames will provide more information about the cell features right before cell division occurs, resulting in increased accuracy of predictions.

**Figure 4:**
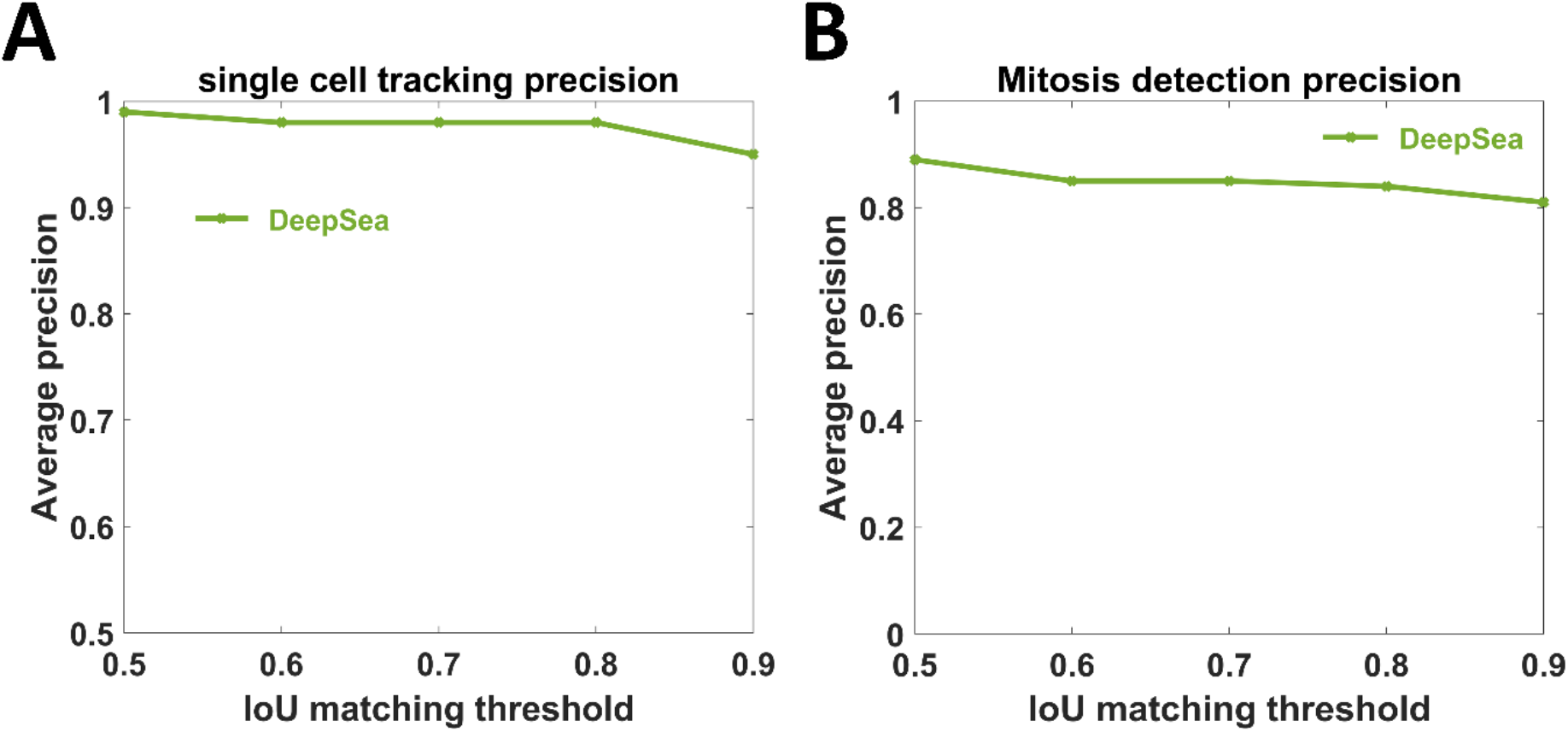
Tracking model evaluation on the test set. We evaluated the model performance using the standard average precision at different IoU matching thresholds. **A**) Single-cell tracking precision at different IoU matching thresholds. **B**) Mitosis detection precision at different IoU matching thresholds.

In another evaluation approach, we assessed our tracker model in a full cell cycle tracking task. This test uses the trained model to track the target cell motion trajectories across the live-cell microscopy frame sequences from birth to division. To be able to test the accuracy of our tracking model, we used MOTA (Multi-Object Tracking Accuracy) which is a widely used metrics in multi-object tracking schemes [20, 21] and measures the precision of localizing objects over time (Equation 5). Our tracker model achieved a MOTA value of 90% (Table 1). Our evaluation process involved 137 full ground truth cell cycle trajectories. Figure 5 shows one example of our model output with MOTA=1.0, tracking the cell motion trajectories over nine consecutive frames for three target cells. We also added the manual editing option to DeepSea software so that users can edit the output of the software and reduce the segmentation and tracking errors to allow researchers to fully track the life cycle of a cell.

**Table 1:** Multi-cell cycle tracking results.

| MOTA | Number of cell cycles | Number of detected cells |
| --- | --- | --- |
| 0.90 | 137 | 2576 |

**Figure 5:**
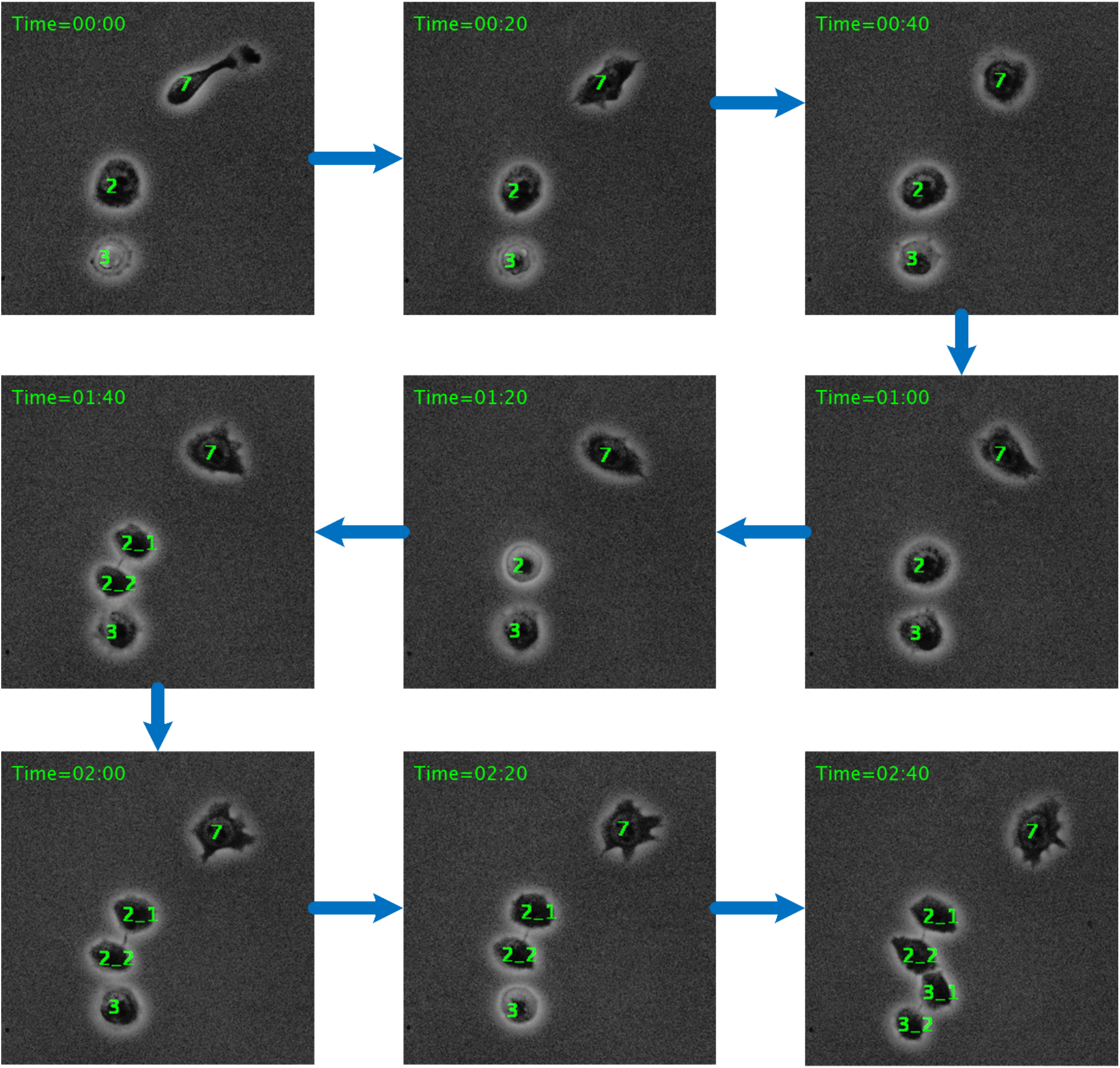
Example of the cell cycle tracking process obtained by feeding nine consecutive stem cell frames (with a sampling time of 20 minutes) to our trained tracking model. Daughter cells are linked to their mother cells by an underline (in the sixth and seventh frames).

### Cell size regulation of mouse embryonic stem cells

Cells need to grow in size before they can undergo division. Different cell types maintain a fairly uniform size distribution by actively controlling their size in the G1 phase of the cell cycle [22]. However, the typical G1 control mechanisms of somatic cells are altered in mESCs [23, 24]. Mouse embryonic stem cells have an unusually rapid cell division cycle that takes about 10h to be completed (Figure 6A). The rapid cell cycle of mESCs is primarily due to an ultrafast G1 phase that is about 2h compared to ~10 in skin fibroblast cells with daughter cells born at different sizes (Supplementary Figure 6A,B). An interesting question is whether mESCs can employ size control in their rapid G1 phase, just as most somatic cells do.

**Figure 6:**
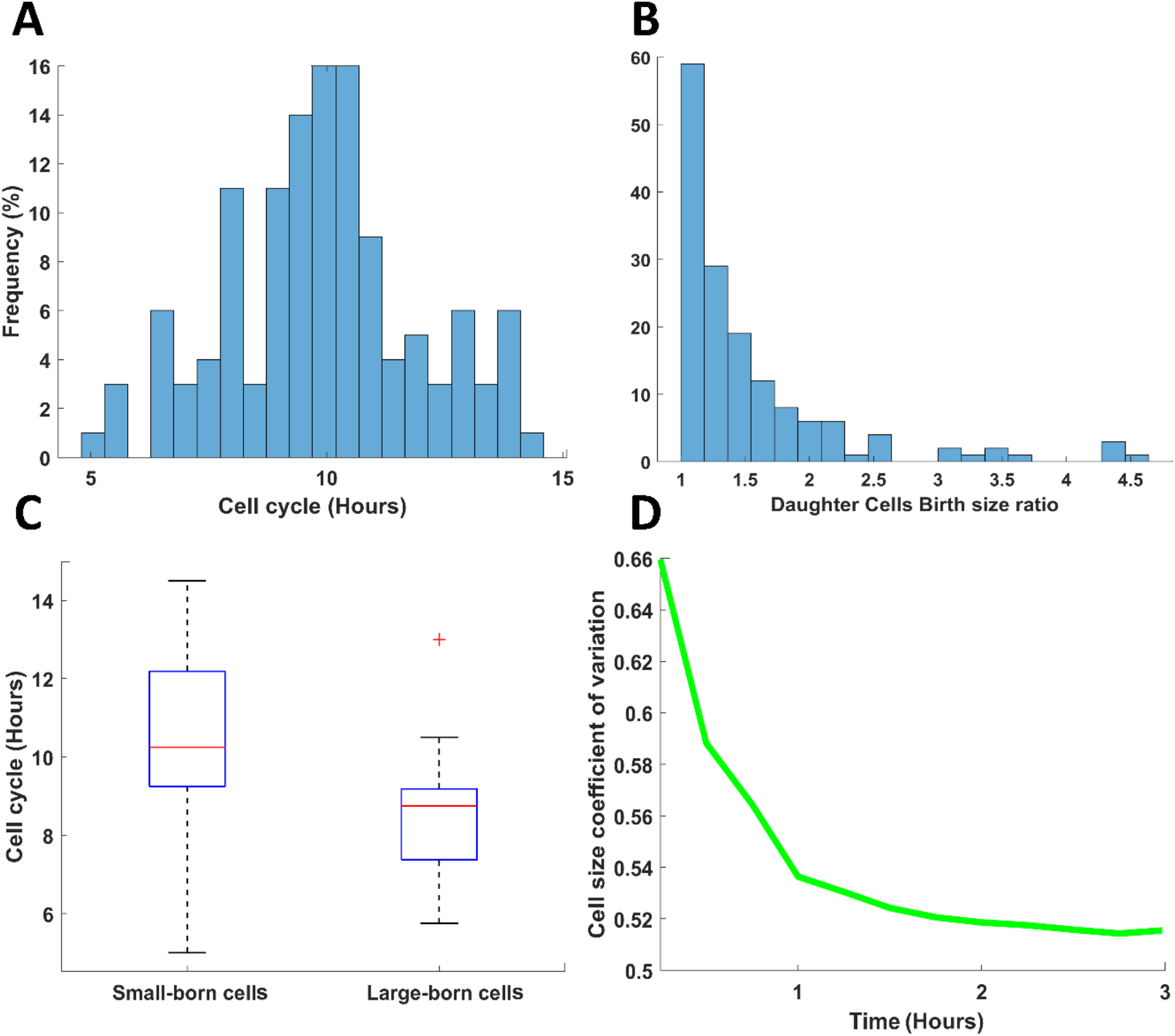
Cell size regulation in mouse embryonic stem cells. **A**) Distribution of the cell cycles. **B**) Histogram of birth size ratio of daughter cell pairs. **C**) Comparing the cell cycle duration of the cells born small with those born large. **D**) Cell size coefficient of variation.

To answer this question, we used our trained DeepSea model to analyze cell size control in mESCs. Using confocal microscopy, we showed that the area of a cell is closely correlated with the cell volume, making the area a faithful measurement for cell size (Supplementary Figure 7). By measuring the size of the sister cells at birth, we showed that 42% of divisions resulted in daughter cells of different sizes (Figure 6B). We hypothesized that smaller born cells would spend more time growing compared to their larger sister cells. In support of this hypothesis, we observed that smaller born cells increase their cell cycle duration by about ~2h compared to their larger sister cells (Figure 6C). Next, we asked if the variations in cell size at birth will be corrected over the course of one cell division cycle by measuring the coefficient of variation (CV) of cell size throughout the cell cycle (Figure 6D). We observed a rapid decline in size variation within the first three hours after birth supporting the hypothesis that mESCs regulate their size in the G1 phase of the cell cycle. Together, our results support the hypothesis that mESCs can adjust the cell cycle based on birth size, suggesting cell size control through an unknown molecular mechanism [22].

## Discussion

Here we introduced DeepSea, a deep learning model for automated analysis of time-lapse images of cells and their nuclei. The segmentation and tracking of cell bodies and subcellular organelles from microscopy images are critical steps for nearly all microscopy-based biological analysis applications. Although phase-contrast microscopy is a non-invasive and widely used method for live-cell imaging, developing automated segmentation and tracking algorithms remains challenging. This is due to low-quality imaging systems, the unpredictable nature of cells in their movements over time, and the proximity of cells. Here, we leveraged the recent advancement in deep learning-based image processing to address some of these challenges.

The lack of a comprehensive, high-quality annotated dataset of cells prevents the full utilization of deep learning-based models for microscopy image analysis systems. We generated large manually annotated datasets of time-lapse microscopy images of three cell types, which are publicly available and can be used for new image analysis models. In addition, we were able to significantly increase the size of annotated data covering more variations by applying image augmentation techniques, which benefited from both conventional image augmentation techniques and a proposed random cell movements method. We expect this resource to facilitate the future application of deep learning-based models for the analysis of microscopy images.

To address the challenge of cell segmentation and tracking, we built a deep learning model, termed DeepSea, which can efficiently segment cell areas in phase-contrast microscopy images. Our segmentation model was trained on our generated dataset and achieved an IOU value of 92% at IOU matching threshold of 0.5. Most importantly, we were able to exploit the deep learning capabilities to automate the tracking of cells across the time-lapse microscopy image sequences. Our DeepSea tracker model was able to track the full cell cycle trajectories with a MOTA value of 90% obtained from 2576 detected cells and 137 cell cycles.

We showcase the application of the DeepSea by investigating cell size regulation in mESCs across hundreds of cell division cycles. Our cell size analysis revealed that smaller born mESCs regulate their size by spending more time growing in the G1 phase of the cell cycle. These findings strongly support the idea that mESCs actively monitor their size, consistent with the presence of size control mechanisms in the short G1 phase of the embryonic cell cycle.

We would like to note that our dataset and models are limited to the phase contrast 2D images of three cell types. However, researcher can train our model using their own annotated images of single cells using DeepSea training options. A larger dataset of the samples from different cell types and different imaging modalities would be useful for testing our proposed model’s generalization, reliability, and robustness. In addition, in our future work, we will investigate other deep models that have recently achieved considerable advancement in object detection and tracking tasks, such as Recurrent Yolo, TrackR-CNN, JDE, RetinaNet, CenterPoint [6–8], or merge their architecture with our current models to improve the results.

## Method

### Dataset

We collected three cell-type datasets of phase-contrast time-lapse microscopy image sequences, including two in-house datasets of Mouse Embryonic Stem Cells (MESC, 31 sets, 1074 images) and Bronchial epithelial cells (7 sets, 2010 images) and one dataset of Mouse C2C12 Muscle Progenitor Cells (7 sets, 540 images) obtained from an external resource with the cell culture described in [25]. Our collected datasets are publicly available on https://deepseas.org/datasets/. Some examples and dataset statistics are shown in Table 2 and Supplementary Figure 8. We designed an annotation software in MATLAB (https://deepseas.org/software/) to manually create the ground-truth mask image corresponding to each stem cell image. We applied a data augmentation scheme to generate a larger dataset with more variations efficiently and less expensively aiming to train a more generalized model. The image augmentation technique offers a solution to create new annotated samples by modifying existing data in a controlled way using a set of image-based or feature-based transformations. Some of these transformations, usually used in image augmentation tasks, are rotation, flipping, reflection, wrapping, adding noise, color space shifting, scaling, cropping, padding, and brightness shifting [26]. In our image augmentation scheme, in addition to conventional image transformations, we proposed moving the stem cell bodies by the random vectors of (θ,d), where θ is the direction angle between 0 and 360 and d is the displacement in pixels.

**Table 2:** Dataset characteristics.

|  | Resolution | Frame Rate | Num of Sets | Num of Images | Num of Single Cells | Num of Cell Cycles |
| --- | --- | --- | --- | --- | --- | --- |
| **Stem Cells** | 1344x1024 | 15-30 min | 30 | 2010 | 14995 | 115 |
| **Bronchial Cells** | 1244x904 | 5 min | 8 | 1174 | 48027 | 292 |
| **Muscle Cells** | 1392x1040 | 20 min | 9 | 502 | 22080 | 274 |

### Cell Culture and Microscopy

Mouse ESCs (V6.5) were maintained on 0.1% gelatin-coated cell culture dishes in 2i media (Millipore Sigma, SF016-100) supplemented with 100U/ml Penicillin-Streptomycin (Thermo Fisher, 15140122). Cells were passaged every 3-4 days using Accutase (Innovate Cell Technologies, AT104) and seeded at a density of 5,000-10,000 cells/cm2. For live imaging, between 5000 to 10,000 cells were seeded on 35mm dishes with a laminin-coated (Biolamina) 14mm glass microwell (MatTek, P35G-1.5-14-C). Cells were imaged in a chamber at 37C perfused with 5% CO2, a Zeiss AxioVert 200M microscope with an automated stage, and an EC Plan-Neofluar 5x/0.16NA Ph1objective or an A-plan 10x/0.25NA Ph1 objective. The same culture condition was used for confocal imaging, except that 24 hours after seeding, the media was replaced with 2ml DMEM-F12 (Thermo Fisher, 11039047) containing 2ul CellTracker Green CMFDA dye (Thermo Fisher, C2925) and placed back in the incubator for 35 minutes. Next, 2 ul of CellMask Orange plasma membrane stain (Thermo Fisher, C10045) was added, and the dish was incubated for another 10 minutes. Dishes were washed three times with DMEM-F12, after which 2ml of fresh 2i media was added. Cells were imaged directly after the live-cell staining protocol using the Zeiss 880 Microscope using a 20x/0.4 N.A. objective and a 1μm interval through the z-axis.

Immortalized human bronchial epithelial (HBEC3kt) cell line homozygous for wildtype *U2AF1* at the endogenous locus was obtained as a gift from the laboratory of Harold Varmus (Cancer Biology Section, Cancer Genetics Branch, National Human Genome Research Institute, Bethesda, United States of America and Department of Medicine, Meyer Cancer Center, Weill Cornell Medicine, New York, United States of America) and cultured according to [27]. This host cell line was used for lentiviral transduction and blasticidin selection to generate a line with stable expression of *KRAS*^G12V^ using a lentiviral plasmid obtained as a gift from the laboratory of John D Minna (Hamon Center for Therapeutic Oncology Research, The University of Texas Southwestern Medical Center) described in [28]. Cells from passage 11 were grown to 80% confluency in Keratinocyte SFM (1X) (Thermo Fisher Scientific, USA) before being re-seeded as biological duplicates at three densities: 0.3M, 0.2M, and 0.5M cells per well in 6-well plates and allowed to adhere before live-cell imaging over a 48h time period.

### Segmentation model

In the instance segmentation task, we proposed and built a 2D deep learning-based model called DeepSea (Supplementary Figure 1). Our model employs residual blocks aiming to(i) increase the depth of the network resulting in fewer extra parameters instead of widening the network, (ii) accelerate the speed of training of the deep networks, (iii) reduce the effect of the vanishing Gradient Problem and (iv) To potentially obtain higher accuracy in network performance as observed in Residual Networks [29]. Our DeepSea model involves only 1.9 million parameters that it is considerably smaller than typical instance segmentation models such as UNET [30], PSPNET [31], SEGNET [32]. We also took advantage of batch normalization and dropout techniques to improve the model’s speed, performance, and stability. The auxiliary touching cell edge representations (highlighting the edge area between touched cells) and the auxiliary training loss value we involved (Equation 2) lead the learning algorithm to spend more computational budget and time on touching cell edges enhance learning edge detection during the training process. To create each touching cell edge mask, we first create a weight map from the ground truth cell masks using the equation 1:

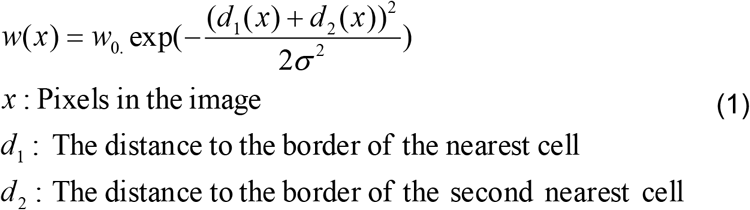

where w_0_ and σ were set to 10 and 25, respectively. Then we make a binary image by replacing all pixel values above a determined threshold (=1.0) with 1s and setting all other pixels to 0s.

In the training process, we used the early stopping technique to stop training when the validation score stops improving. We used 60% of the data samples for the training process, and thus the remaining data were used in the validation and test process equally to evaluate the trained model’s performance. We chose the RMSprop optimization function with the learning rate scheduler of the OneCycleLR method (LR=1e-3) to optimize model weights and minimize the proposed loss function (Equation 2). Our loss function is a linear combination of cross-entropy (CE) loss and Dice loss (DL) functions [33] as well as auxiliary loss functions (EdgeCE and EdgeDL) for the touching cell edge representations. CE takes care of pixel-wise prediction accuracy, while DL helps the learning algorithm increase the overlap between true area and predicted area, which is essentially needed where the number of image background pixels is much higher than foreground pixels (object area pixels).

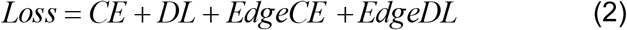

In the test phase, we used the IOU index, a value between 0 and 1 and known as the Jaccard index as well [34] (Equation 3), to match the segmentation model predictions to the ground truth annotated masks:

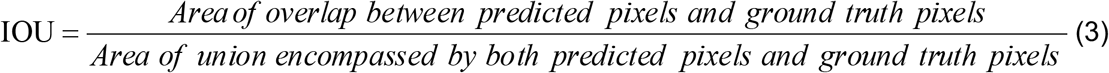

In each test image, we labeled each detected cell body whose IOU index was higher than a predefined threshold value as a valid match and so True Positive (TP) prediction. Also, the ground truth cell body masks with no valid match were categorized into the False Negative (FN) set, and the predictions with no valid ground truth masks were labeled as the False Positive (FP) cases (non-cell objects). Then using Equation (4), we calculated the average precision (AP) value for each image in the test set, used by the other state of the art methods in cell body segmentation tasks [16]:

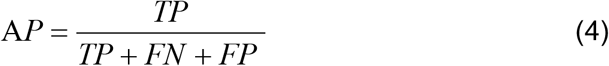

### Tracking model

Our tracking model aims to detect the same cell bodies from one frame to the next and detect cell divisions (mitosis). The cell tracking process helps create cell lineage structure over time. We used an architecture baseline like the DeepSea segmentation model but with multiple images, two inputs, and one output (Supplementary Figure 3). The first input is the target cell image at the previous time point and the second input is the segmented cell image at the current time point, and the output is the binary cell body mask of the target cell at the current time point. This model extracts the features of the single target cell in the previous frame to localize and detect it (or its daughter cells) among the segmented cells in the current frame (Supplementary Figure 4). To increase the accuracy of the tracking model, we limited our search space (at time t) in x and y coordinates to a small square with the size of 6 times the target body size) centered at the previous frame cell’s centroids (at time t-τ). Since cells move slowly through space, the cell’s previous location present a good guess of where the model should expect to find it in the current frame.

The number of the model parameter is only 2.1 million, while the other most used tracking models, such as ROLO [35], DeepSort [36], TrackRCNN [37], use more than 20 million parameters, confirming that we have an efficient model in the tracking process as well. We used image augmentation techniques during the training process to increase the train data variations and make a more generalized and stronger model. Also, since the number of cell division events is naturally much less than single-cell tracking events, the tracking model potentially tends to overfit single-cell tracking samples. To reduce the risk of overfitting, we artificially repeated the cell division events fifty times more in our train dataset using image augmentation techniques to help the model see a balanced number of single-cell tracking events and cell division events during the training process. The train optimization function and hyperparameters are the same as the segmentation model training process.

In the test phase, we used the same IOU metric (Equation 3) to match the binary masks generated by the tracking model to the ground truth annotated masks. We also used the AP introduced in Equation (4) to measure the average precision of the model in tracking the cell bodies from one frame to the next and detecting cell divisions. To evaluate our tracking model in a continuous cell trajectory tracking process during an entire cell life cycle from birth to division, we used MOTA (Multiple Object Tracking Accuracy, Equation 5), widely used in multi-object tracking challenges [20, 21]. To our knowledge, this is the first time that this metric has been used to evaluate a cell tracking model performance.

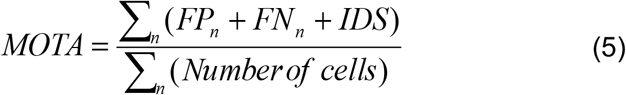

where n is the frame number and IDS (Identity Switches) are the number of times a cell is assigned a new label during its life cycle. A perfect tracker model achieves MOTA=1.

### Software availability

All implemented methods are provided as Python scripts and can be downloaded from https://github.com/abzargar/DeepSea. Also, our DeepSea software is available on https://deepseas.org/software/ with examples and instructions. DeepSea is a user-friendly software designed to enable researchers to 1) load and explore their phase-contrast cell images in a high contrast display, 2) detect and localize cell bodies, 3) track and label cell lineages across the frame sequences, 4) manually correct the DeepSea models’ outputs, 5) train a new model with a new cell type dataset, 6) save the results and reports on the local system. It employs our last trained DeepSea models in the segmentation and tracking processes.

## Author Contribution

A.Z.K and S.A.S conceived the project and wrote the manuscript. S.A.S performed live microscopy imaging. A.Z.K developed the DeepSea software and implemented the machine learning code. A.B and E.H.R contributed to the generation bronchial epithelial cell images. A.Z.K, N.C, N.M, C.W.N and K.A contributed to the generation of a manually annotated dataset of images. G.L and N.C performed user quality control of DeepSea segmentation and tracking functions. G.L, S.R and L.H contributed to the confocal microscopy images and provided feedback to improve our manuscript.

## Acknowledgment

This work was supported by the NIGMS/NIH through a Pathway to Independence Award K99GM126027/ R00GM126027 (S.A.S.), start-up package of the University of California, Santa Cruz (S.A.S), R01HD098722 (LH) and James H. Gilliam Fellowships for Advanced Study program (SR). We acknowledge core support from the UCSC Institute for the Biology of Stem Cells (IBSC), IBCS’ imaging facility (SCR_021135), CIRM Shared Stem Cell Facilities (CL1-00506-1,2) and CIRM Major Facilities (FA1-00617-1).

## References

[1] J. Fiorentino, M. Torres, and A. Scialdone, “Measuring and Modeling Single-Cell Heterogeneity and Fate Decision in Mouse Embryos,” Annual Review of Genetics, Vol. 54, 2020.

[2] Bogdan P. et al., “Heterogeneous Structure of Stem Cells Dynamics: Statistical Models and Quantitative Predictions,” Nature Scientific Reports, Vol. 4, 2014.

[3] Semrau S. et al., “Dynamics of lineage commitment revealed by single-cell transcriptomics of differentiating embryonic stem cells,” Nature Communications, Vol. 8, 2017.

[4] Stavroula Skylaki, Oliver Hilsenbeck & Timm Schroeder, “Challenges in long-term imaging and quantification of single-cell dynamics,” Nature Biotechnology, Vol. 34, 2016.

[5] Anatole Chessel; Rafael E. Carazo Salas, “From observing to predicting single-cell structure and function with high-throughput/high-content microscopy,” Vol. 63, No. 2, pp. 197–208, 2019.

[6] Zatulovskiy, E. et al., “Cell growth dilutes the cell cycle inhibitor Rb to trigger cell division,” Science, Vol. 369, pp. 466–471, 2020

[7] Skylaki, S. et al., “Challenges in long-term imaging and quantification of single-cell dynamics,” Nat Biotechnol, Vol. 34, pp. 1137–1144, 2016.

[8] G. Ciaparrone et al., “Deep learning in video multi-object tracking: A survey,” Neurocomputing, Vol. 381, pp. 61–88, 2020.

[9] S. Yun and S. Kim, “Recurrent YOLO and LSTM-based IR single pedestrian tracking,” IEEE Conference on Control, Automation and Systems, pp. 15–18, 2019.

[10] X. Zhou, D. Wang, P. Krähenbühl, “Objects as Points,” arXiv:1904.07850, 2019.

[11] Ouyang, W. et al. Deep learning massively accelerates super-resolution localization microscopy. Nat. Biotechnol. 36, 460–468 (2018).

[12] Wang, H. et al. Deep learning enables cross-modality super-resolution in fluorescence microscopy. Nat. Methods 16, 103–(2019).

[13] Beier, T. et al. Multicut brings automated neurite segmentation closer to human performance. Nat. Methods 14, 101–102 (2017).

[14] Weigert, M. et al. Content-aware image restoration: pushing the limits of fluorescence microscopy. Nat. Methods, 15, 1090–1097 (2018).

[15] Wu, Y. et al. Three-dimensional virtual refocusing of fluorescence microscopy images using deep learning. Nat. Methods, 16, 1323–1331 (2019).

[16] Stringer, C., Wang, T., Michaelos, M. et al. Cellpose: a generalist algorithm for cellular segmentation. Nat Methods 18, 100–106 (2021).

[17] Schmidt, U., Weigert, M., Broaddus, C. & Myers, G. Cell detection with star-convex polygons. International Conference on Medical Image Computing and Computer-Assisted Intervention 265–273 (2018)

[18] Liu, L. et al., “Deep Learning for Generic Object Detection: A Survey,” International Journal of Computer Vision, Vol. 128, pp. 261–318, 2020.

[19] Minaee, S. et al., “Image Segmentation Using Deep Learning: A Survey,” arXiv:2001.05566v5, 2020.

[20] Ciaparrone, G. et al., “Deep Learning in Video Multi-Object Tracking: A Survey,” Computer Vision and Pattern Recognition, arXiv:1907.12740, 2019.

[21] Ristani, E. et al., “Performance Measures and a Data Set for Multi-Target, Multi-Camera Tracking,” Computer Vision and Pattern Recognition, arXiv:1609.01775, 2016.

[22] Evgeny Zatulovskiy and Jan M. Skotheim, “On the Molecular Mechanisms Regulating Animal Cell Size Homeostasis,” Trends Genetics, Vol. 36, pp. 360–372, 2020.

[23] Ben Boward, Tianming Wu, and Stephen Dalton, “Concise Review: Control of Cell Fate Through Cell Cycle and Pluripotency Networks,” Stem Cells, Vol. 34, pp. 1427–1436, 2016.

[24] Liu, L. et al., “The cell cycle in stem cell proliferation, pluripotency and differentiation,” Nat Cell Biol, Vol. 21, pp. 1060–1067, 2019.

[25] KER, DAI FEI ELMER. “Phase Contrast Time Lapse Microscopy Image Datasets with Human-Generated Ground Truths and Computer-Aided Cell Tracking Annotations.” OSF, 9 July 2020. Web.

[26] Connor Shorten & Taghi M. Khoshgoftaar, “A survey on Image Data Augmentation for Deep Learning,” Journal of Big Data, Vol. 6, No. 60, 2019.

[27] Fei, D. L. et al.,“Wild-Type U2AF1 Antagonizes the Splicing Program Characteristic of U2AF1-Mutant Tumors and Is Required for Cell Survival,” PLoS Genet, Vol. 12, pp. 1–26, 2016.

[28] Sato, M. et al., “Human lung epithelial cells progressed to malignancy through specific oncogenic manipulations,” Mol Cancer Res., Vol. 11, pp. 638–50, 2013.

[29] Khan, A. et al., “A survey of the recent architectures of deep convolutional neural networks,” Artificial Intelligence Review, Vol. 53, pp. 5455–5516, 2020.

[30] Olaf Ronneberger, Philipp Fischer, Thomas Brox, “U-Net: Convolutional Networks for Biomedical Image Segmentation,” Computer Vision and Pattern Recognition, arXiv:1505.04597, 2015.

[31] Zhao, H. et al., “Pyramid Scene Parsing Network,” Computer Vision and Pattern Recognition, arXiv:1612.01105, 2017.

[32] Vijay Badrinarayanan, Alex Kendall, Roberto Cipolla, “SegNet: A Deep Convolutional Encoder-Decoder Architecture for Image Segmentation,” Computer Vision and Pattern Recognition, arXiv:1511.00561, 2016.

[33] Shruti Jadon, “A survey of loss functions for semantic segmentation,” Image and Video Processing, arXiv:2006.14822, 2020.

[34] Rezatofighi, H. et al., “Generalized Intersection over Union: A Metric and A Loss for Bounding Box Regression.” Computer Vision and Pattern Recognition, arXiv:1902.09630, 2019.

[35] Ning, G. et al, “Spatially Supervised Recurrent Convolutional Neural Networks for Visual Object Tracking”, Computer Vision and Pattern Recognition, arXiv:1607.05781, 2016.

[36] Wojke, N. et al., “Simple Online and Realtime Tracking with a Deep Association Metric,” Computer Vision and Pattern Recognition, arXiv:1703.07402, 2017.

[37] P. Voigtlaender et al., “MOTS: Multi-Object Tracking and Segmentation,” IEEE Conference on Computer Vision and Pattern Recognition (CVPR), 2019.

