## Supplementary Figures for "DeepSea: An efficient deep learning model for single-cell segmentation and tracking of time-lapse microscopy images"

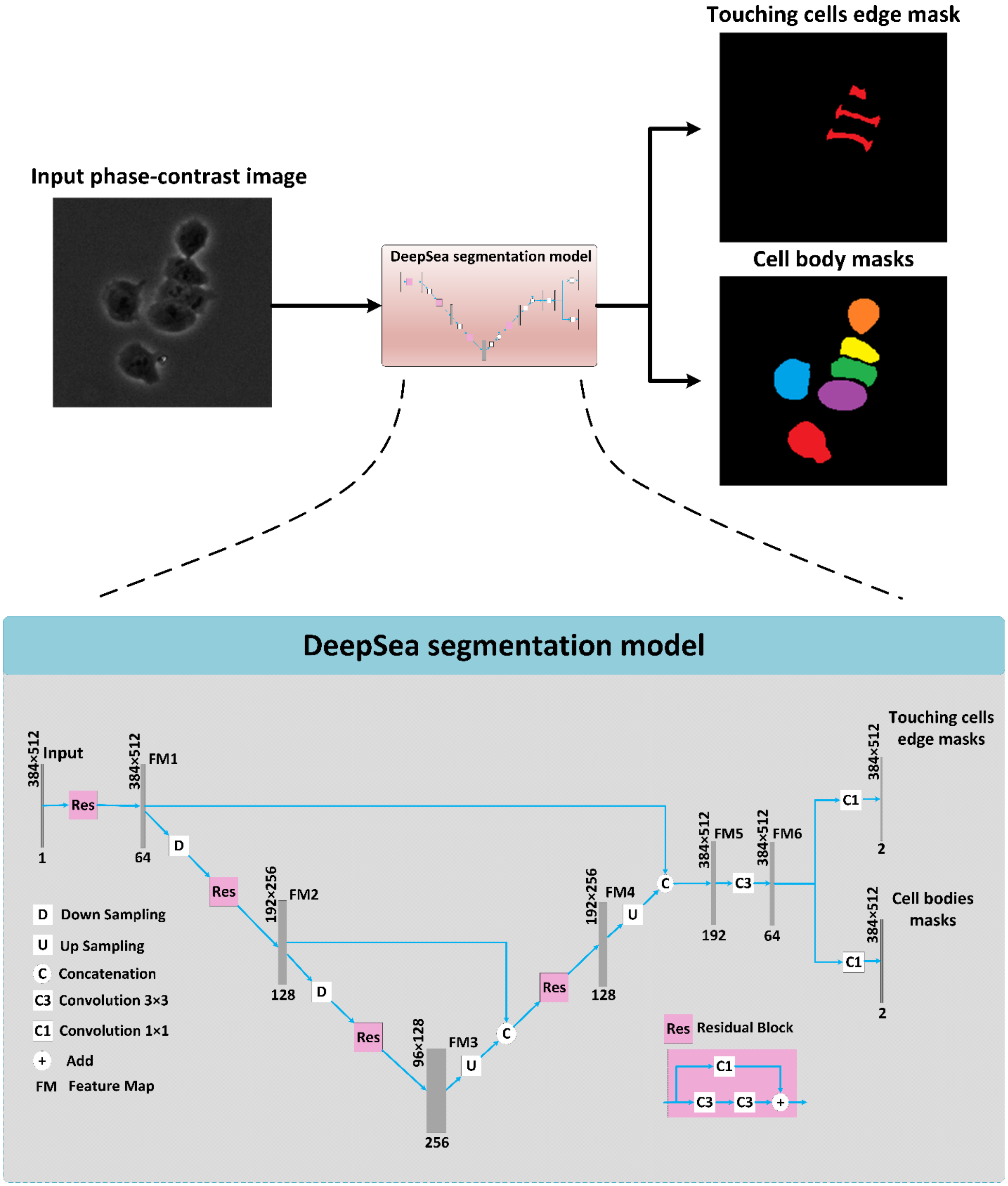


**Supplementary Figure 1**: Our DeepSea segmentation model uses residual blocks to make the segmentation mor efficient and the auxiliary touching cell edge representations helps to enhance the performance of the model in high density cultures.

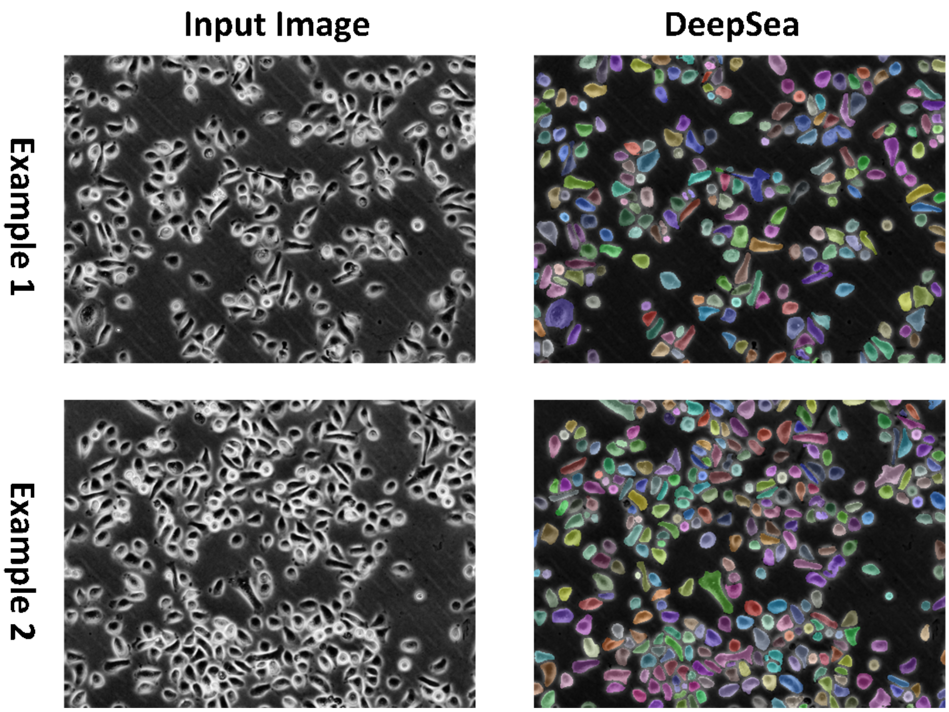


**Supplementary Figure 2**: Two examples of the DeepSea segmentation model output showing high accuracy with high-density cell images.

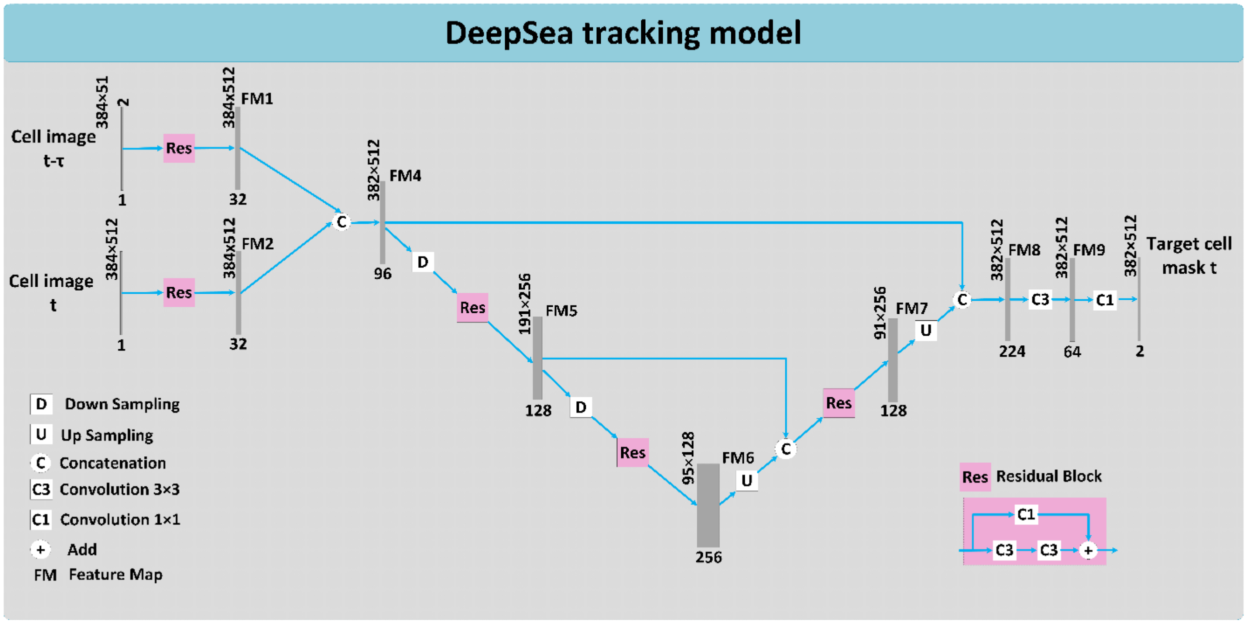


**Supplementary Figure 3**: DeepSea tracking model architecture with two input images of subsequent timepoints and the output of target cell mask.

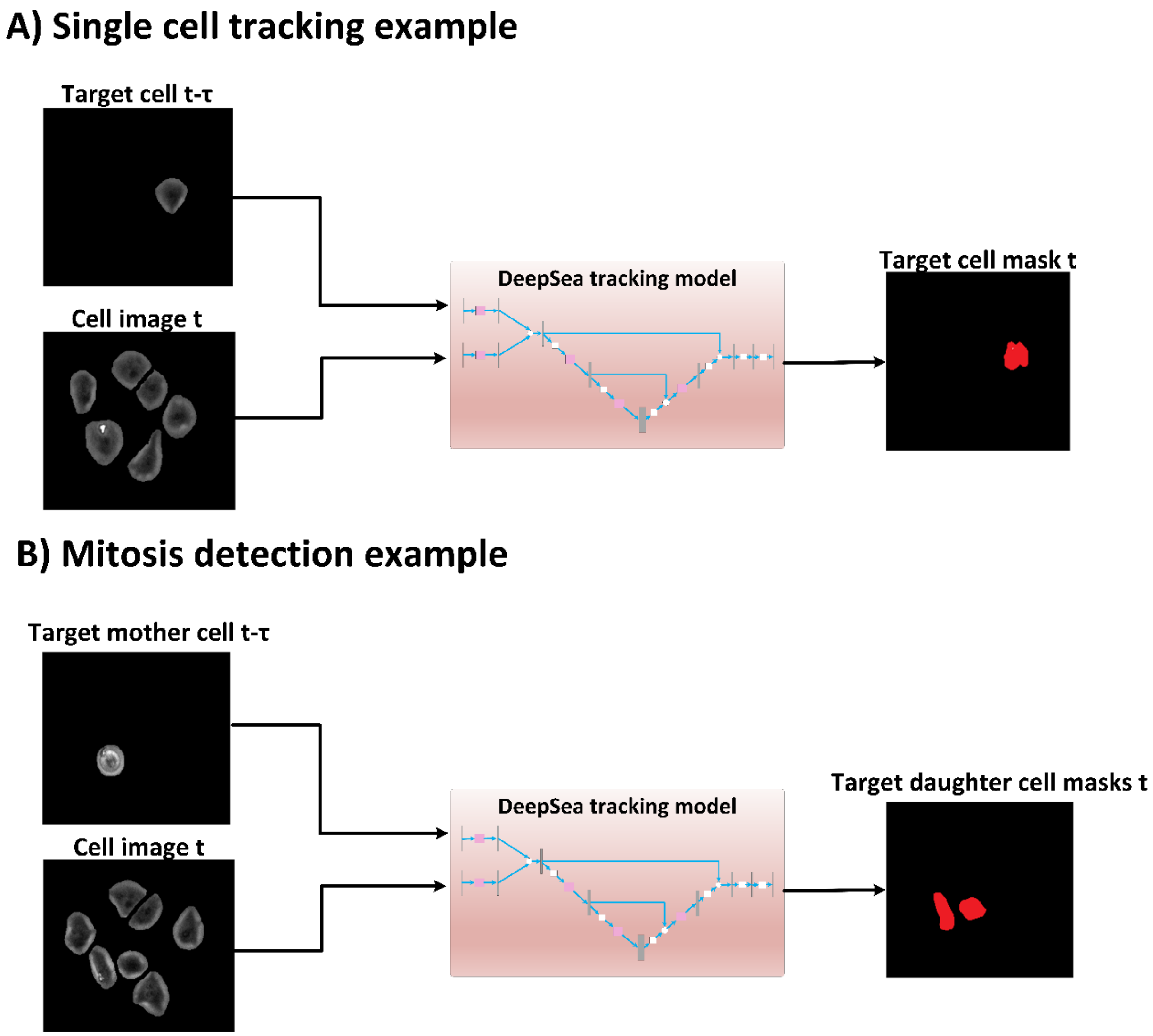


**Supplementary Figure 4**: **A**) Single-cell tracking example from one frame to the next frame. **B**) Daughter cell detection example from one frame to the next frame.

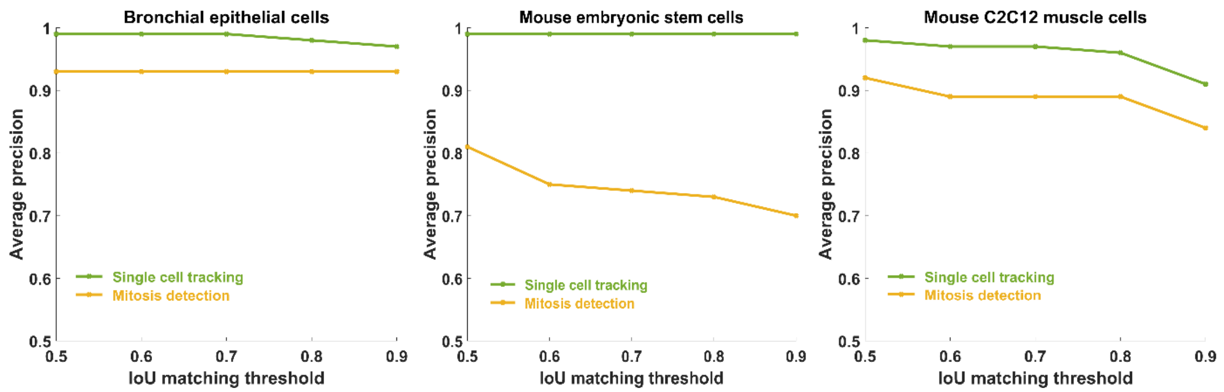


**Supplementary Figure 5**: The DeepSea tracker model and mitotic detection performance with different cell type test sets.

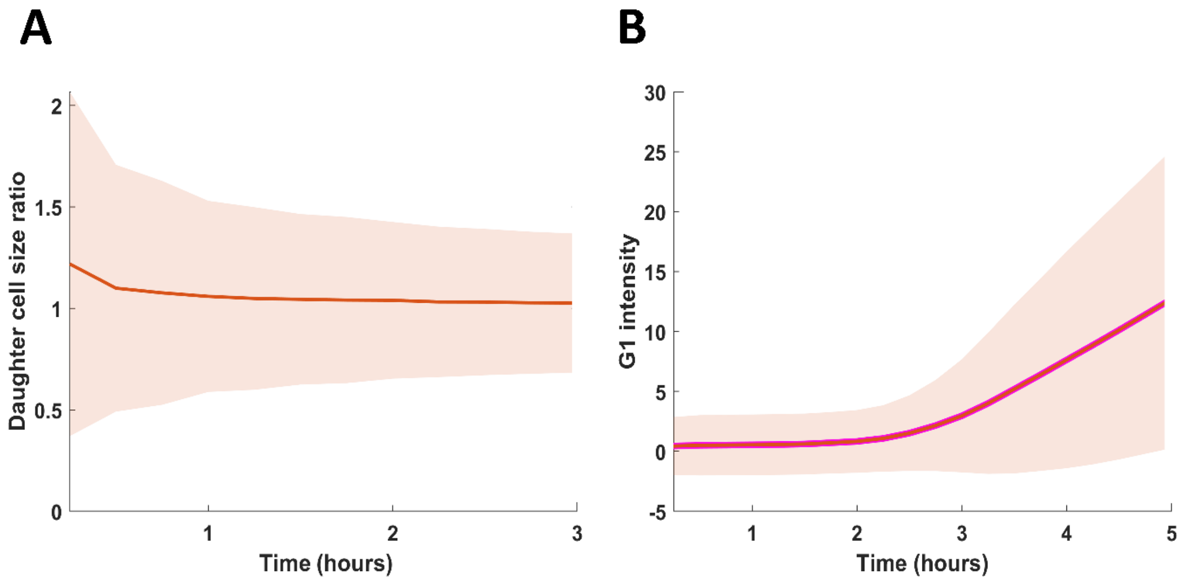


**Supplementary Figure 6**: **A**) Cell size ratio graph of daughter cell pairs. **B**) Automated measurement of G1 duration using Fucci sensor.

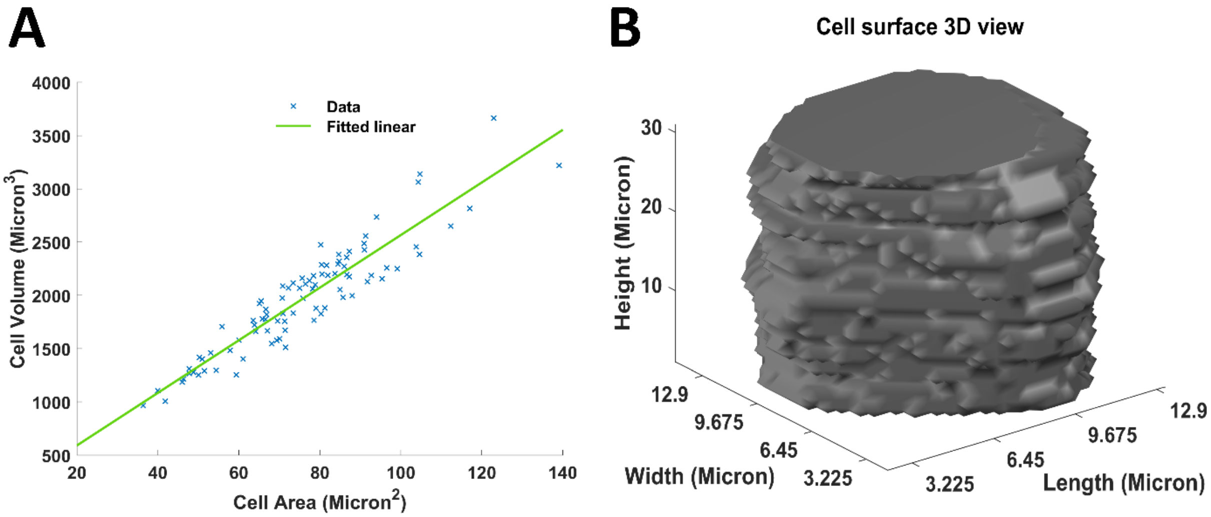


**Supplementary Figure 7**: **A**) Cell area versus cell volume measurement using confocal microscopy for each embryonic stem cell. **B**) One example of cell surface measurement in our confocal microscopy experiment.

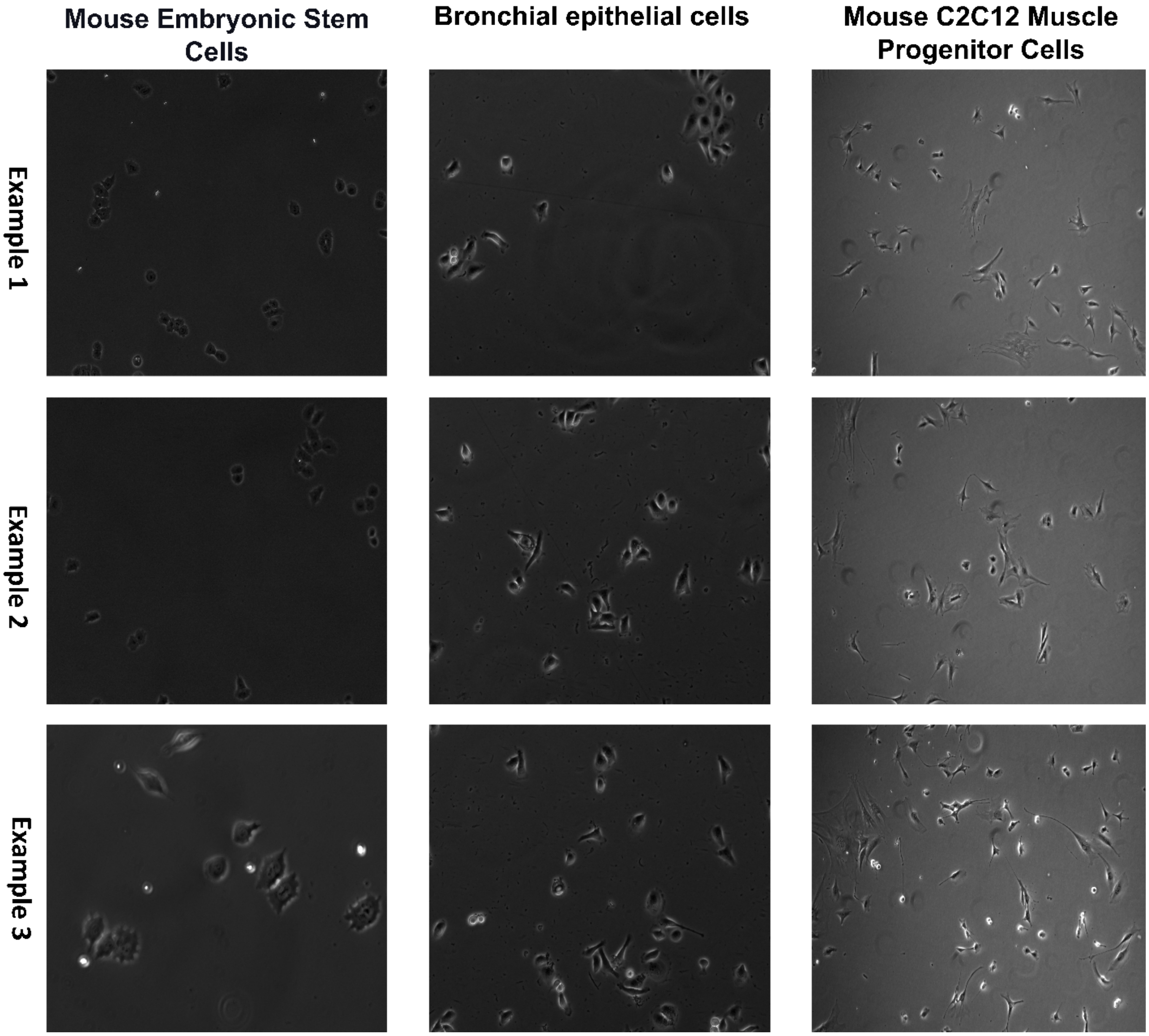


**Supplementary Figure 8**: Examples of three cell types used in our dataset, including Mouse Embryonic Stem Cells, Bronchial epithelial cells, and Mouse C2C12 Muscle Progenitor Cells.
